# Orthogonal spectral and temporal envelope representation in auditory cortex

**DOI:** 10.1101/2024.10.28.620417

**Authors:** Kuniyuki Takahashi, Tianrui Guo, Tatsuya Yamagishi, Shinsuke Ohshima, Hiroaki Tsukano, Arata Horii

**Affiliations:** Department of Otolaryngology Head and Neck Surgery, Faculty of Medicine, University of Miyazaki, Miyazaki 889-1692, Japan; Department of Otolaryngology, Graduate School of Medicine and Dental Sciences, Niigata University, Niigata 951-8510, Japan; Department of Psychiatry, University of North Carolina at Chapel Hill, Chapel Hill, NC 27599, USA; Neuroscience Center, University of North Carolina at Chapel Hill, Chapel Hill, NC 27599, USA

## Abstract

Speech perception relies on two fundamental acoustic components: spectral and temporal. While spectral information is known to be represented in the auditory cortex through tonotopy, how temporal features are organized has remained unclear. Here, by varying rise-ramp steepness and frequencies, we reveal that the steepness of the temporal envelope—a critical cue for phonemes discrimination and sound source perception—is systematically mapped in the mouse auditory cortex. Using widefield calcium imaging, we discovered that the envelope steepness is represented orthogonally to the tonotopic axis, forming a two-dimensional cortical map that mirrors the dual structure of sounds. This organization was observed in primary-like auditory regions but not in higher-order-like areas, indicating distinct auditory processing streams. These findings uncover a principle of cortical organization, suggesting that the auditory cortex encodes sound along two independent axes and thereby provides a neural basis for parallel processing for complex sounds such as speech and natural acoustic environments.

## Introduction

Speech sounds are acoustically composed of two main components: spectral and temporal (Figure 1A). Spectral components provide cues critical for speech perception: pitch cues, which support intonation and sex recognition, and spectral-shape cues, where differences in overall frequency distribution and formant structure—shaped by the place of articulation—serve as acoustic cues that listeners use to identify speakers (Oganian et al., 2023; Shannon et al., 1995). Temporal components, particularly the sound envelope that reflects dynamic amplitude fluctuation over time (Drullman, 1995), also play a critical role in speech perception (Rosen, 1992). Multiple lines of evidence support this: speech signals with similar envelope shapes yield similar percepts, enabling accurate phoneme recognition even across speakers with different fundamental frequencies (Aizawa and Eggermont, 2006; Frye et al., 2007). Psychological studies using synthesized sounds with selectively emphasized components have shown that the envelope provides more robust cues for speech perception than the carrier frequency (Smith et al., 2002), and high speech recognition can be achieved with temporal cues alone, even in the absence of detailed spectral information (Shannon et al., 1995). Since the temporal envelope is a conserved feature of vocalizations across mammalian species (Figure 1B), the spectral–temporal dichotomy likely supports effective vocal communication across a broad range of species, and their cortical representations offers insight into general principles of auditory coding.

**Figure 1.**
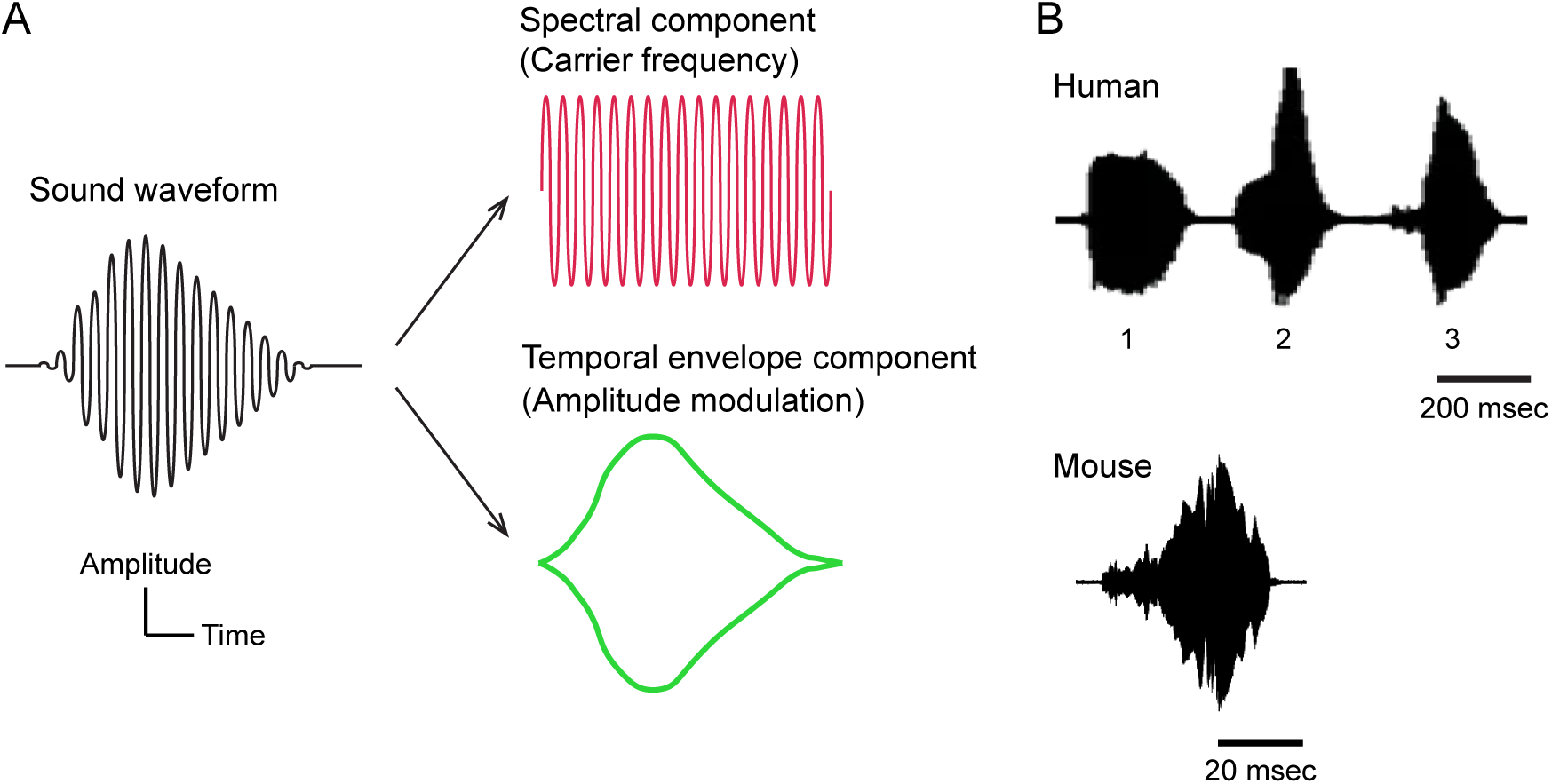
Spectral and temporal components of sound waveforms. (A) Two principal acoustic components of sound: spectral and temporal. Spectral components correspond to the rapid oscillations of sound waves, also referred to as carrier frequencies or temporal fine structures. They are important for pitch perception and for conveying formant frequencies that underlie the perception of vocal sounds. Temporal components reflect the slower fluctuation in amplitude over time, commonly known as the temporal envelope, which plays a key role in the perception of voiced sounds. The perception of voiced phonemes relies on the integration of both spectral and temporal cues. **(B)** Example waveforms of vocalized sounds of mammals. The top waveform shows human speech. Waveforms labeled 1, 2, and 3 correspond to the phonemes /go/, /le/, and /ku/, respectively (Milne et al., 2016). Rise time is distinct for different phenomes; for example, it is 8 msec for /ba/ and 45 msec for /wa/ (Nittrouer et al., 2013). The bottom waveform illustrates a mouse vocalization. Both human and mouse examples demonstrate clear temporal envelopes, a feature conserved across mammalian vocalizations. The top images are adapted from Milne et al. (2016) under a Creative Commons license. The bottom image reuses data from Aponte et al. (2021).

Functional organization in the cortex is a fundamental principle of mammalian sensory processing, and the auditory cortex exhibits a prominent topographic organization. It is well-established that the auditory cortex exhibits tonotopy, in which neurons are spatially arranged according to preferred frequency (Bizley et al., 2005), and causal manipulations have demonstrated that activation in specific locations drives corresponding percepts (Ceballo et al., 2019). In contrast, much less is known whether envelopes are represented, although it is already known that envelope perception is mediated by auditory cortical activity from clinical cases involving damage to the human auditory cortex (Maffei et al., 2007). Animal studies using single-neuron recordings have revealed temporal envelope sensitivity in auditory neurons (Eggermont, 2001; Fishbach et al., 2001; Heil, 2001; Nagarajan et al., 2002; Phillips et al., 2002). Yet how these local responses are integrated into larger-scale cortical organization has remained unresolved. Absence of clear evidence may stem from technical limitations of sparse recordings, which lack the spatial resolution needed to map out potential fine functional organizations.

To overcome this gap, we characterized the cortical representation of temporal envelopes using widefield calcium imaging in GCaMP6f-expressing mice. This technique enables fine-scale mapping of cortical functional organizations with its high spatial resolution (Issa et al., 2014; Romero et al., 2020; Calhoun et al., 2023). Our investigation focused on the steepness of the envelope, a key feature in shaping the envelope, and sought to determine the presence or absence of a functional organization related to the envelope. We employed simplified acoustic stimuli that systematically varied onset ramp steepness, a paradigm used in many previous physiological investigations (Suga, 1971; Phillips, 1998; Heil, 2001; Peterson and Heil, 2021). This stimulus is particularly advantageous for GCaMP macroscale imaging, as GCaMP reports clear responses during the onset phase of sound presentation, with less sustained activity during the ongoing phase (Issa et al., 2014; Romero et al., 2020; Chen et al., 2021). We discovered that two primary-like auditory regions exhibit a two-dimensional map, in which envelope steepness is represented orthogonally to tonotopy. In contrast, higher-order-like regions represent only tonotopy. Together, these findings reveal a dual-axis principle of cortical representation, which aligns two fundamental acoustic dimensions—frequency and temporal envelope—within the mouse auditory cortex, providing a neural substrate for perception of complex sounds such as speech and natural acoustic environments.

## Results

### Visualizing auditory cortex maps with calcium imaging

Calcium signals were simultaneously recorded from the bilateral auditory cortices of GCaMP6f-expressing mice, with our initial analysis focused on the right auditory cortex (Figure 2A and 2B). To visualize the tonotopic maps, mice were exposed to tones with frequencies ranging from 5 to 60 kHz, following the parameters used in our previous studies (Takahashi et al., 2006; Tsukano et al., 2015). Fluorescent changes were consistently observed in response to the onset of tones across all frequencies (Figure 2C). Although various naming conventions exist for areas of the mouse auditory cortex, in this study, we refer to auditory areas according to the classification presented by Tsukano et al., 2017. Clear tonotopic gradients were observed in the anterior auditory field (AAF), dorsal medial field (DM), primary auditory field (A1), and secondary auditory field (A2), and the directions of tonotopic gradients were consistent with many previous studies (Figure 2D and 2E) (Tsukano et al., 2016; Liu et al., 2019; Narayanan et al., 2022). Although the presence of other areas such as the dorsoanterior (DA), dorsoposterior (DP), or ventroposterior auditory field (VPAF) have been reported, we did not evaluate them in the present study, because these regions do not exhibit clearly localized responses to pure tone stimuli in C57BL/6-based mice, therefore their tonotopy cannot be reliably visualized.

**Figure 2.**
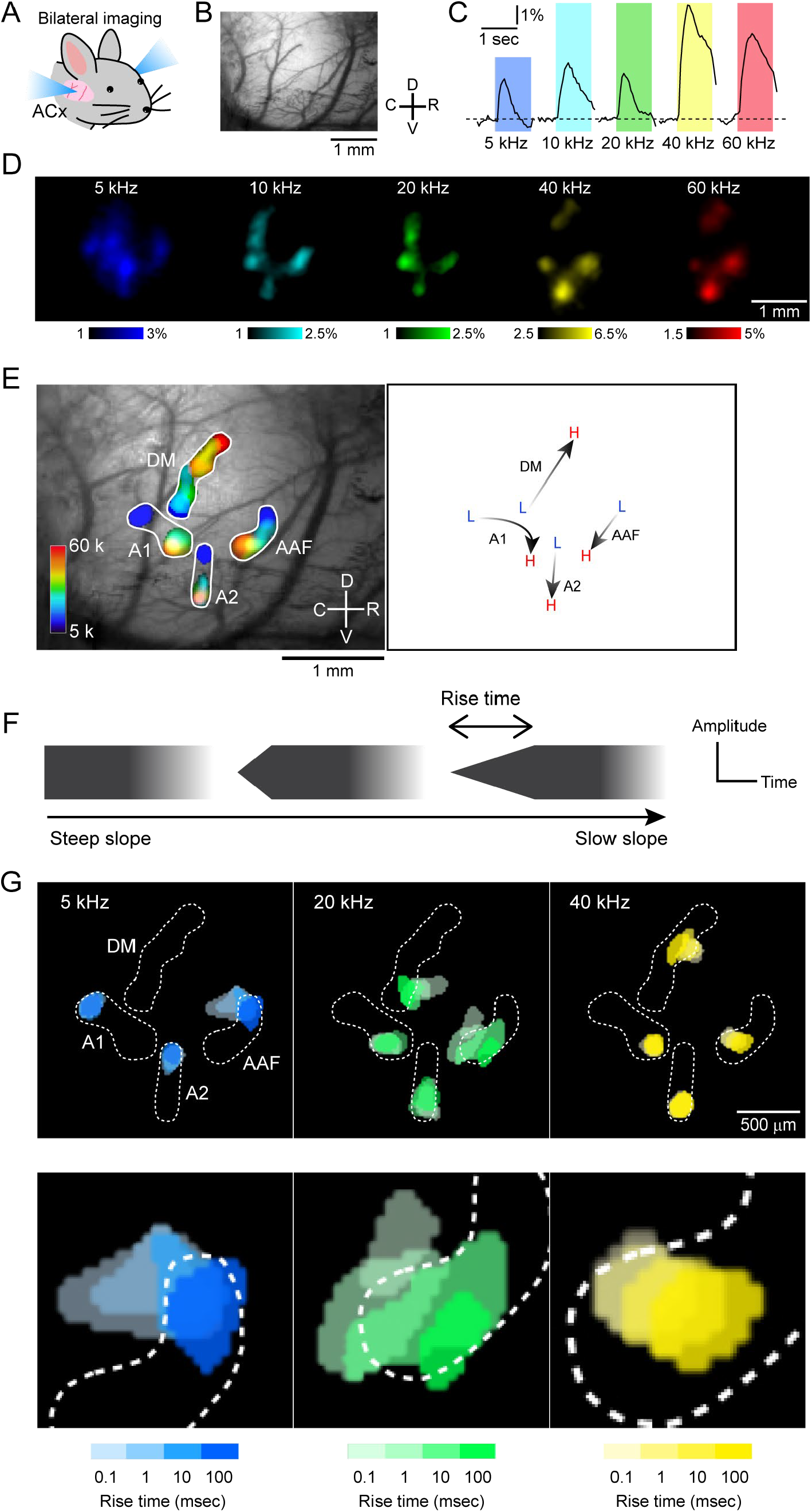
Visualizing envelope steepness maps orthogonal to tonotopy in right auditory cortex. **(A)**Illustration of bilateral auditory cortex imaging. ACx, auditory cortex. **(B)** An example field of view of the right auditory cortex. C, caudal; D, dorsal; R, rostral; V, ventral. **(C)** Example temporal profiles of GCaMP6f signals in AAF in response to tones with frequencies of 5–60 kHz. Tonal presentation periods were shown in color. **(D)** Example images of peak responses for 5–60 kHz tones. Color bar values indicate ΔF/F_0_ (%). **(E)** Left: Tonotopic maps of the right auditory cortex visualized by superimposing thresholded response images shown in (D). AAF, anterior auditory field; A1, primary auditory field; A2, secondary auditory field; DM, dorsomedial field. Right: Low to high tonotopic directions of each auditory area in the example mouse. **(F)** Schematic explaining the change in rise ramp of tonal onset. **(G)** Example maps of rise-ramp steepness, obtained from the same mouse as above. Thresholded peak responses visualized using tones with rise-ramp times ranging from 0.1 to 100 msec were superimposed. The gradation in color indicates rise times of 0.1, 1, 10, and 100 msec from light to dark. Areas enclosed by white dots indicate the boundaries of tonotopic maps shown in (E). Magnified AAF images were shown at the bottom.

In order to evaluate the impact of the temporal envelope steepness on the auditory cortex, we employed simplified stimuli that systematically varied onset ramp steepness, following previous studies (Phillips, 1998; Heil, 2001). We varied the rise-ramp steepness of tones presented to the same mouse, with ramp durations ranging from 0.01 to 100 msec, and carrier frequencies of 5, 20, and 40 kHz (Figure 2F and 2G). Surprisingly, we observed a clear shift in the peak location of tonal responses especially in AAF and DM, occurring in a direction orthogonal to the tonotopic gradient.

### Presence of two-dimensional map in primary-like regions

To examine these maps in greater detail, we performed a quantitative analysis of group data from AAF. In the 2-D plots, tonotopic organization is seen in the dorso-ventral direction (Figure 3A), while the shorter rise time, i.e., steeper rise ramp, shifted peak response locations in the rostro-caudal direction regardless of carrier frequency (Figure 3A). Notably, tonotopic organization remained the same across different rise ramp steepness. These findings suggest that AAF represents a two-dimensional map on the cortical surface. The mean angle between the tonotopy and rise-ramp map was approximately 82 degrees (Figure 3B), suggesting that these two maps were distinct and orthogonal to each other. A statistically significant correlation was found between rise-ramp steepness and the distance orthogonal to tonotopy (r = 0.63, p = 5.8 × 10^-10^; Figure 3C, left). It is worthwhile to note that the regression line aligns well on the logarithmic-scale horizontal axis for rise-ramp times, which suggests that the variations in steepness, particularly when the onset ramp is steep, are captured with high spatial resolution. In contrast, no correlation was observed between rise-ramp steepness and the direction along tonotopy (r = -0.064, p = 0.57; Figure 3C, right).

**Figure 3.**
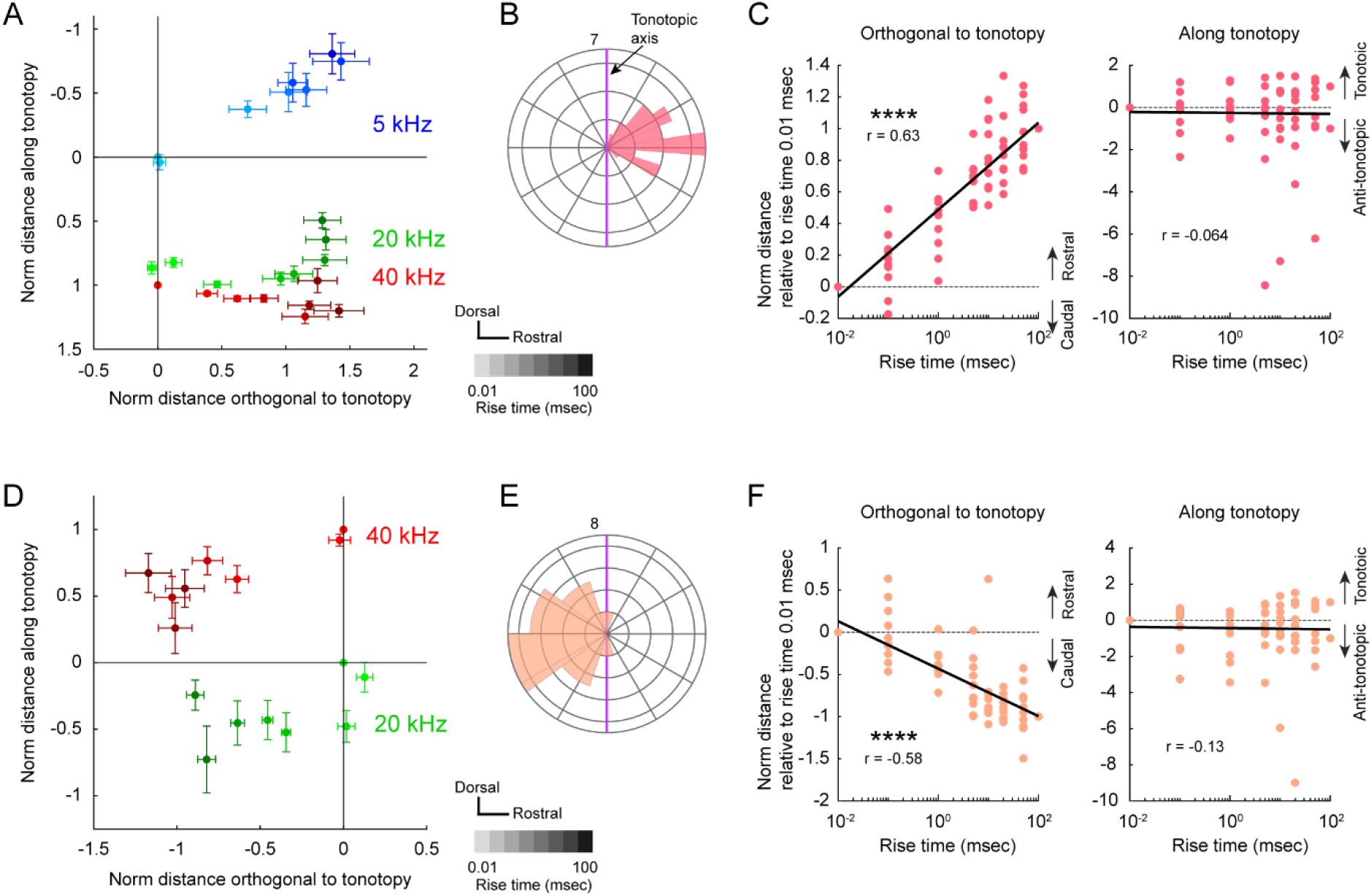
Presence of envelope maps in AAF and DM. **(A)** Group data showing the location of response peaks for tones with various rise-ramp times in AAF. The response location is evaluated using 5, 20, and 40 kHz tones with rise-ramp times ranging from 0.01 to 100 msec. The gradation in color indicates rise rimes of 0.01, 0.1, 1, 5, 10, 20, 50, and 100 msec from light to dark. Coordinates are normalized using the distance between peaks for 5 kHz and 40 kHz tones with a rise-ramp time of 0.01 msec, applied to both horizontal and vertical axes. Data are presented in mean ± SEM. **(B)** Circular histogram showing the direction of responses of rise-ramp times of 100 msec relative to those for rise-ramp times of 0.01 msec. Purple lines indicate the tonotopic axis. Data for 5, 20, 40 kHz are co-plotted. Orientation is consistent to (A). n = 30 plots from 10 mice. **(C)** Left: Relationship between rise-ramp time and the shift relative to the peak location for a rise-ramp time of 0.01 msec in the direction orthogonal to tonotopy in AAF. r = 0.63, ****p = 5.8 × 10^-10^ (Pearson’s correlation); n = 80 plots from 10 mice. Right: Relationship between rise-ramp time and the shift relative to the peak location for rise-ramp time of 0.01 msec in the tonotopic direction. r = -0.064, p = 0.57. **(D)** Same as (A) but for DM. Coordinates are normalized using the distance between peaks for 20 kHz and 40 kHz tones with a rise-ramp time of 0.01 msec for horizontal and vertical axes. n = 30 plots from 10 mice. **(E)** Same as (B) but for DM. **(F)** Same as (C) but for DM. Right, r = -0.58, ****p = 2.0 × 10^-8^. Left, r = -0.13, p = 0.26; n = 80 plots from 10 mice.

Similarly, analysis of DM revealed a rise-ramp map that was a mirror image of that in AAF (Figure 3D–F). Due to the instability of low-frequency responses in DM, we conducted a quantitative analysis of DM maps, excluding the 5 kHz responses. The peak response locations shifted caudo-rostrally with increasing rise ramp steepness, irrespective of carrier frequency (Figure 3D). The angle between the tonotopic map and the rise-ramp map in DM was approximately 74 degrees (Figure 3E). A statistically significant correlation was also observed between rise-ramp steepness and the distance orthogonal to tonotopy in DM (r = 0.58, p = 2.0 × 10^-8^; Figure 3F, left), while no correlation was found between rise-ramp steepness and the direction along tonotopy (r = 0.13, p = 0.26; Figure 3F, right). In summary, both AAF and DM exhibit mirror-imaged rise-ramp maps that are orthogonal to each area’s tonotopic organization.

We conducted similar analyses in A1 and A2. Interestingly, neither A1 nor A2 exhibited detectable rise-ramp maps. In A1, tonotopic organization runs in the caudo-rostral direction, curving in the high frequency part (Figure 4A), as reported previously (Tsukano et al., 2015). Interestingly, there was no shift in response peaks perpendicular to the tonotopic axis, even when the rise-ramp steepness was altered (Figure 4A and 4B). No statistically significant correlation was observed between rise-ramp steepness and the distance orthogonal to tonotopy (r = 0.0052, p = 0.96; Figure 4C, left), and there was no significant correlation between rise-ramp steepness and the direction along tonotopy (r = 0.21, p = 0.059; Figure 4C, right). Similarly, in A2, altering the rise-ramp steepness did not cause any shift in response peaks perpendicular to the tonotopic axis (Figure 4D and 4E). No statistically significant correlation was found between rise-ramp steepness and the distance orthogonal to tonotopy (r = -0.053, p = 0.64; Figure 4F, left), and there was no significant correlation between rise-ramp steepness and the direction along tonotopy (r = 0.044, p = 0.70; Figure 4F, right). These results suggest that A1 and A2 primarily represent one-dimensional maps reflecting tonotopy, without distinct representation of rise-ramp steepness.

**Figure 4.**
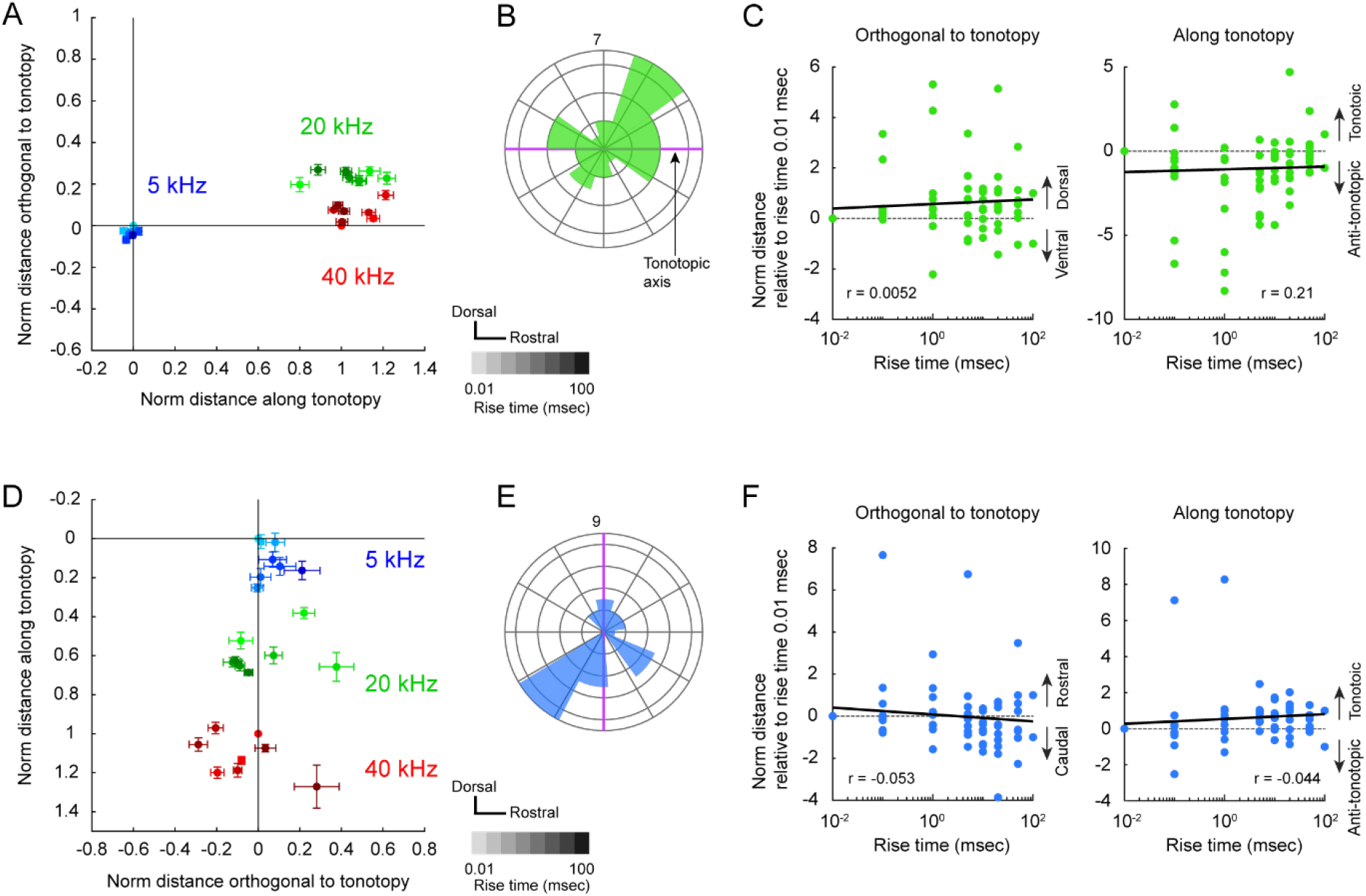
Absence of envelope maps in A1 and A2. **(A)** Group data showing the location of response peaks for tones with various rise-ramp times in A1. The response location is evaluated using 5, 20, and 40 kHz tones with rise-ramp times ranging from 0.01 to 100 msec. The gradation in color indicates rise times of 0.01, 0.1, 1, 5, 10, 20, 50, 100 msec from light to dark. Coordinates are normalized using the distance between peaks for 5 kHz and 40 kHz tones with a rise-ramp time of 0.01 msec, applied to both horizontal and vertical axes. Data are presented in mean ± SEM. **(B)** Circular histogram showing the direction of responses of rise-ramp times of 100 msec relative to those for rise-ramp times of 0.01 msec. Purple lines indicate the tonotopic axis. Data for 5, 20, 40 kHz are co-plotted. Orientation is consistent to (A). n = 30 plots from 10 mice. **(C)** Left: Relationship between rise-ramp time and the shift from the peak location for a rise-ramp time of 0.01 msec in the direction orthogonal to tonotopy in A1. r = 0.0052, p = 0.96 (Pearson’s correlation); n = 80 plots from 10 mice. Right: Relationship between rise-ramp time and location shift relative to the peak location for rise-ramp time of 0.01 msec in the tonotopic direction. r = 0.21, p = 0.059. **(D)** Same as (A) but for A2. n = 29 plots from 10 mice. **(E)** Same as (B) but for A2. **(F)** Same as (C) but for A2. Right, r = -0.053, p = 0.64. Left, r = 0.044, p = 0.70; n = 77 plots from 10 mice.

### Bilateral representation of two-dimensional map

We performed simultaneous imaging of the bilateral auditory cortex in the same group of mice, allowing us to examine whether the rise-ramp map is represented in both hemispheres. We found similar functional organizations in the left auditory cortex. As rise-ramp steepness increases, the responses shifted caudally in AAF, while the responses in DM shifted rostrally (Figure S1A), as seen in the right hemisphere. On the other hand, clear shifts were not seen in A1 or A2. A statistically significant correlation was found between rise-ramp steepness and the distance orthogonal to tonotopy in AAF (r = 0.60, ****p = 3.9 × 10^-9^) and DM (r = -0.40 ***p= 2.1 × 10^-4^), but not in A1 (r = 0.026, p = 0.83) or A2 (r = -0.11, p = 0.33) (Figure S1B). No correlation was found between rise-ramp steepness and the direction along tonotopy in any areas (AAF, r = 0.076, p = 0.50; DM, r = 0.15, p = 0.19; A1, r = 0.012, p = 0.92; A2, r = 0.11, p = 0.35) (Figure S1C). These data suggest that the AAF and DM symmetrically represent rise-ramp maps across hemispheres.

### Map shift purely caused by change in rise steepness

It is well known that the spectrum of pure tones becomes splattered when the rise time is extremely short, such that additional surrounding frequencies are emitted from the speaker in addition to the expected pure tone (Hartmann and Sartor, 1991). Although this effect is evident for rise times shorter than 1 msec in our on-site experience, it has been little investigated in recording studies. To examine this in our setup, we recorded pure tones with rise times ranging from 0.01 to 100 ms and generated spectrograms. Narrow, symmetrical spectral splatter was detected around the carrier frequency in 5-kHz tones with rise times of 0.01–1 ms (gray arrowheads, Figure 5A). Unexpectedly, a broader splatter with a lower-frequency bias was observed at 0.01 and 0.1 ms (red arrowheads, Figure 5A,B). However, clear splatter was not observed for 5 kHz tones with rise times longer than 10 ms, nor under any rise-time condition at 20 or 40 kHz (Figure S2C). These results indicate that the gradual orthogonal map shifts observed at 20 and 40 kHz cannot be explained by spectral splatter.

**Figure 5.**
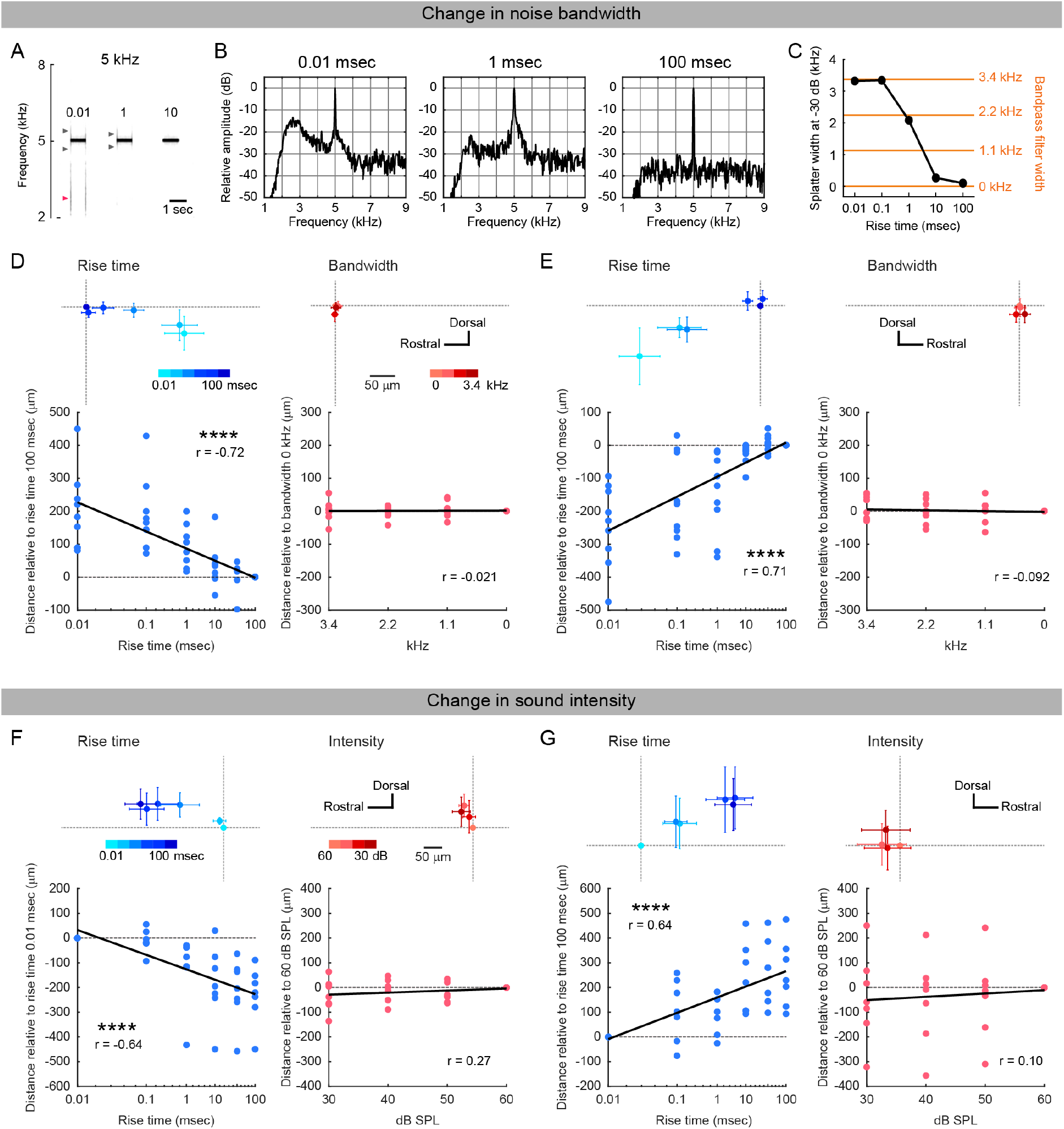
Absence of map shifts caused by acoustic features other than rise steepness. (A) Spectrogram of 5 kHz tones with rise times of 0.01, 1, 100 msec. Gray arrowheads indicate small, symmetrical spectral splatter around the carrier frequency, and red arrowheads indicate broader, lower-frequency–biased splatter. Spectrograms of other frequencies are shown in Figure S2A. **(B)** Amplitude spectra of 5 kHz tones with rise times of 0.01, 1, and 100 msec at tone onset. Y-axis values are expressed in dB relative to the peak amplitude at the carrier frequency. Amplitude spectra of all tones are shown in Figure S2B. **(C)** Splatter width of 5 kHz tones across rise times. Width was quantified as the -30 dB span of Gaussian fits (see also Figure S2). Yellow values indicate the width of band noise added to 5 kHz. **(D)** Left: Relationship between rise-ramp time and location shift relative to the peak location for 5 kHz with a 100 ms rise time in left AAF. Top: group data shown in mean ± SEM. Bottom: correlation along the axis orthogonal to tonotopy. r = -0.72, ****p = 9.9 × 10^-10^ (Pearson’s correlation); n = 54 plots from 9 mice. Right: Relationship between band-noise width and location shift relative to the 0-width condition in left AAF. r = 0.021, p = 0.90; n = 36 plots from 9 mice. Two maps were obtained from the same field of view. **(E)** Same as (D), but for right AAF. Rise time: r = 0.71, ****p = 1.9 × 10^-9^; n = 54 plots from 9 mice. Band noise width: r = -0.092, p = 0.60; n = 36 plots from 9 mice. **(F)** Left: Relationship between rise-ramp time and location shift relative to the peak location for 5 kHz with a 0.01 msec rise time in left AAF. Top, group data. Bottom, correlation along the axis orthogonal to tonotopy. r = -0.64, ****p = 4.7 × 10^-6^; n = 42 plots from 7 mice. Right: Relationship between band-noise width and location shift relative to the 0-width condition. r = 0.22, p = 0.27; n = 28 plots from 7 mice. Two maps were obtained from the same field of view. **(G)** Same as (F) but for right AAF. Rise time: r = 0.64, ****p = 5.7 × 10^-6^; n = 42 plots from 7 mice. Band-noise width: r = 0.10, p = 0.60; n = 28 plots from 7 mice.

To further test whether splatter itself could cause orthogonal map shifts, we quantified the width of splatter in 5 kHz tones (Figure S2) and added band-pass noise spanning this frequency range to 5 kHz tones (Figure 5C). We then assessed whether such noise induced orthogonal map shifts in AAF, by comparing response locations to 5 kHz tones with rise times of 0.01–100 ms and 5 kHz tones with band-pass noise from 0–3.4 kHz in the same animals. Band noise was added to 5 kHz tones with a 100 ms rise time, since this condition alone produces no splatter. While robust rostrocaudal maps of rise-ramp steepness were consistently observed in both hemispheres (left, r = -0.72, ****p = 9.9 × 10^-10^; right, r = 0.71, ****p = 1.9 × 10^-9^) (Figure 5D,E), adding band noise did not induce spatial shifts (left, r = 0.021, p = 0.90; right, r = -0.092, p = 0.60) (Figure 5D,E). These findings suggest that rise-ramp steepness, rather than spectral splatter associated with short rise times, is the principal parameter underlying orthogonal map shifts.

Because shallower onset steepness could also reduce overall sound intensity, we next tested whether intensity changes contributed to orthogonal map shifts. We varied the intensity of 5 kHz tones with a fixed 0.01-msec rise time from 30 to 60 dB SPL and assessed response locations using the same coordinate framework as for the rise-steepness map in the same animals. We used a 0.01-ms rise time to decouple intensity from onset steepness, since adjusting amplitude of tones with long rise times affects onset steepness. We found that response locations in AAF did not shift as a function of sound intensity in either hemisphere (left, r = 0.22, p = 0.27; right, r = 0.10, p = 0.60) (Figure 5F,G). Together with findings from a previous two-photon imaging study (Bandyopadhyay et al., 2010), these results suggest that sound intensity is not functionally organized in the mouse auditory cortex at either the macroscopic or single-neuron level. Finally, we tested whether laterality of sound input could account for map shifts, as early studies suggested that contralateral vs. ipsilateral inputs might be spatially organized within iso-frequency bands in auditory cortex (Middlebrooks et al., 1980). However, tonal response locations were unaffected by the stimulated ear (Figure S3), indicating the absence of a macroscopic laterality map in the mouse auditory cortex, consistent with prior reports (Panniello et al., 2018). Together, these results demonstrate that the orthogonal map in the mouse auditory cortex reflects the steepness of sound onset ramps, but not other acoustic or biological factors such as spectral splatter, sound intensity, or input laterality.

### Orthogonal shifting during the ramping phase

Although we observed rise-ramp steepness-related shifts in response location in AAF and DM, the GCaMP signals were slow, with peak responses occurring after the tonal waveform had reached its plateau phase. To verify that response locations differ during the ramping phase, we visualized responses specifically during the ramping phase by performing imaging at a higher sampling rate (Figure S4). We used tones with rise ramp times of 75 msec and 200 msec and compared the location of peak responses in AAF using data at the timing of 40–60 msec after the tonal onset to ensure evaluation of the rise-ramp map during the ramping phase. The location of peak responses for 200 msec rise-ramp time was significantly more rostral than that for the 75 msec ramp time (p = 0.039, two-sided paired t-test). This data confirms that positional shifts occurred definitely during the ramping phase, suggesting that response location is purely sensitive to rise-ramp steepness. Overall, our study demonstrates that AAF and DM represent two-dimensional maps encompassing both tonotopy and rise-ramp steepness, with the rise-ramp maps displayed in a mirror-imaged fashion and separated by the center area (the dorsoanterior field, DA) (Figure 6).

**Figure 6.**
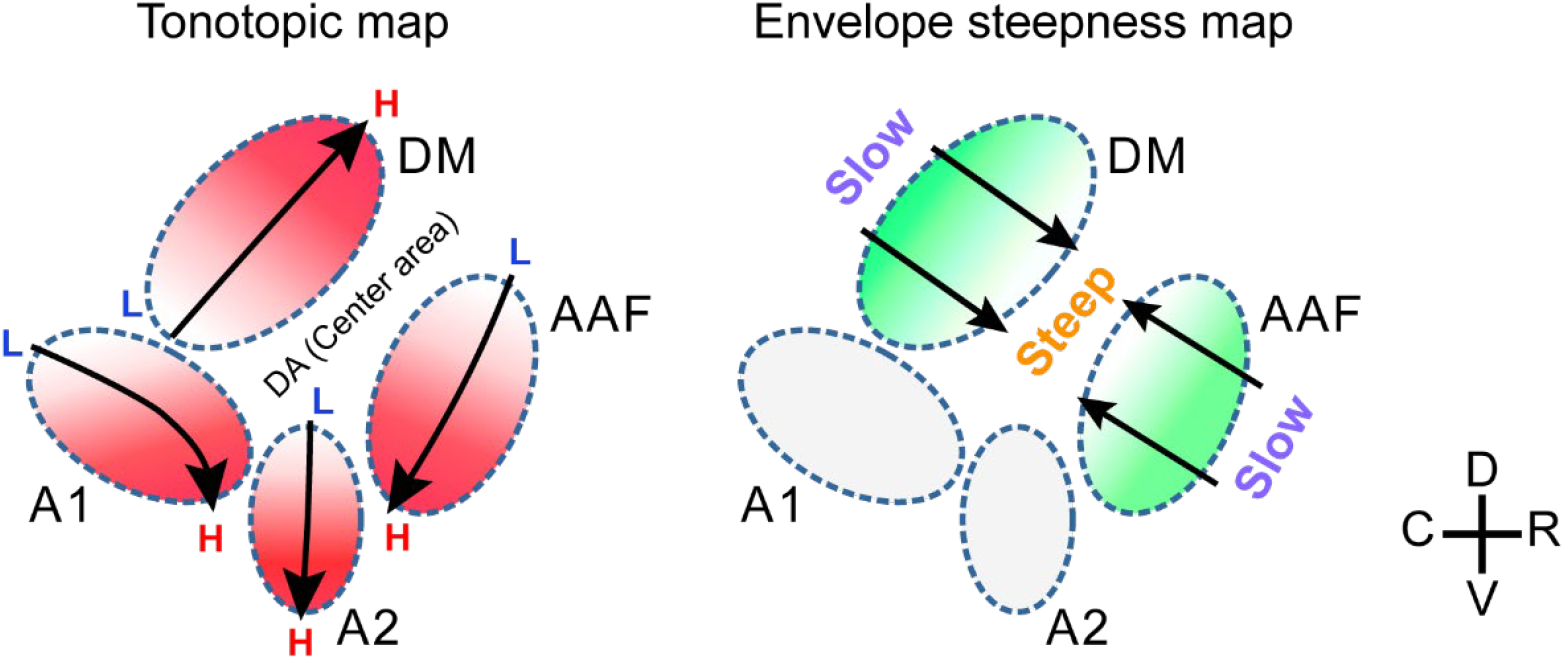
Summary illustration of two-dimensional maps present in mouse auditory cortex. Left: Tonotopic maps in the mouse auditory cortex. The mouse auditory cortex is thought to comprise at least four tonotopic regions, with an additional center area (the dorsoanterior field (DA))(Tsukano et al., 2017) located between AAF and DM. Many studies have shown that the center area does not respond to simple pure tone stimulation (Honma et al., 2013; Issa et al., 2014, 2016; Tsukano et al., 2015; Ceballo et al., 2019; Liu et al., 2019; Narayanan et al., 2022). Right: Envelope steepness maps revealed in the current study. These maps run orthogonal to the tonotopic maps in AAF and DM. In AAF, steepness increases from rostral to caudal, whereas in DM, it increases from caudal to rostral.

## Discussion

The present study reveals that mouse auditory cortex contains well-defined two-dimensional maps where tonotopy and rise-ramp steepness are represented orthogonally. While tonotopy is a prominent feature that is topographically represented in the auditory cortex, the existence of the information encoded in the secondary dimension has been a longstanding question. In the mouse auditory cortex, previous studies have reported functional organizations distinct from tonotopy, such as representation of frequency modulation sweeps (Issa et al., 2016) and offset responses (Baba et al., 2016; Liu et al., 2019); however, these were not represented as gradual topographic gradients independent of tonotopy. In establishing the rise-ramp steepness map, we carefully controlled for confounding factors, as gradual changes in onset steepness inherently affect other acoustic parameters, such as spectral splatter at very short rise times or variations in sound intensity. Yet these factors (Figure 5), as well as input laterality (Figure S3), could not explain the map shifts observed here. Therefore, to the best of our knowledge, this is the first demonstration of two-dimensional maps in the mouse auditory cortex where tonotopy and envelope steepness are orthogonally organized.

Systematic spatial representation of temporal envelope steepness suggests that the mouse auditory cortex encodes selectivity for envelope steepness. This finding advances the understanding of temporal envelope processing, building on earlier electrophysiological studies, which demonstrated neurons consistently responding to the same steepness (Heil, 1997, 1998; Heil and Irvine, 1998) and neurons selective for the direction of envelope slope (Lu et al., 2001; Zhou and Wang, 2010). While the auditory cortex is known to be critical for perceiving both of temporal envelopes (Nourski et al., 2009) and pitch (Warrier and Zatorre, 2004), the orthogonal representation of envelope steepness and tonotopy suggests that temporal envelopes are processed independently of spectral features in the auditory cortex, supporting the psychological observation that temporal envelope and carrier frequency work independently in verbal communication (Smith et al., 2002). Our study underscores a fundamental biological principle: dichotomous functional structures mirror two acoustic features critical for vocalization, linking neural circuitry with acoustic signals of vocal communication. While this principle should apply broadly across mammals, identifying it in the mouse auditory cortex is especially valuable given the increasing focus on vocalization processing in mice. This finding lays the groundwork for exploring how spectral and temporal information are integrated in the auditory cortex, and paves the way toward addressing higher-order questions about how animals perceive conspecific vocalizations (Stiebler et al., 1997; Agarwalla et al., 2023).

Previous studies reported that the orthogonal map to tonotopy represents periodotopy—the cycle of peaks in amplitude-modulated sound waveforms. This has been observed in several species other than mice, throughout different stages of the auditory pathway (Schreiner and Langner, 1988; Krishna and Semple, 2000; Langner et al., 2002, 2009; Baumann et al., 2011, 2015; Barton et al., 2012). In contrast, our findings suggest that the presentation of a one-time rise-ramp steepness, which does not inherently involve periodicity, is sufficient to elucidate an orthogonal map in the mouse auditory cortex. Two possible explanations could reconcile our findings with previous reports. First, the mouse auditory system may differ from that of other animals, with cortical representations being particularly sensitive to envelope steepness. As a result, the mouse auditory cortex developed a specialized functional organization to process this aspect in detail. Second, envelope steepness might be a primary factor in visualizing the orthogonal map in previous studies using other animals. In nearly all previous studies, amplitude-modulated tones were used as periodic acoustic stimuli, which involve repeated ramps with consistent rise and fall steepness, making it difficult to determine whether periodicity or repetitive slope plays a greater role in visualizing the orthogonal map. To determine whether periodicity alone is sufficient to visualize the second map, click trains, which lack envelope ramps, would be an ideal stimulus. However, few studies have used them, and one failed to visualize orderly periodotopy in the inferior colliculus (Schnupp et al., 2015). Future research should disentangle the relative contribution of periodicity and ramp steepness to the visualization of secondary dimensional maps. Due to the temporal limitations of GCaMP signals, the present study primarily focused on the steepness of sound onsets using classical paradigms. Further investigation will be required to determine whether our findings generalize to amplitude fluctuations during the ongoing portion of sounds. Nevertheless, because periodicity and envelopes are sometimes discussed interchangeably, this study highlights the importance of recognizing envelope steepness and periodicity as biologically distinct concepts (Rosen, 1992).

Interestingly, we observed region-specific differences in the presence of envelope steepness maps: clear two-dimensional maps were observed in AAF and DM, whereas such organization was not apparent in A1 and A2. Based on the latency (Guo et al., 2012), the broadness of frequency tuning (Guo et al., 2012; Joachimsthaler et al., 2014), and the connectivity patterns with the auditory thalamus (Ohga et al., 2018), the regions labeled as AAF and DM in the present study exhibit more primary-like characteristics, while A1 appears intermediate, and A2 is considered a higher-order area (Tsukano et al., 2017; Kline et al., 2021). These distinctions may suggest functional specialization within the mouse auditory cortex, with primary-like regions such as AAF and DM preferentially more engaged in processing envelope features. This interpretation is potentially consistent with prior findings from other species, which have demonstrated more robust temporal coding in primary auditory areas than in higher-order auditory areas (Schreiner and Urbas, 1988; Eggermont, 1998). Alternatively, the apparent absence of orthogonal maps in higher-order–like areas may result from transformations of functional organization occurring along the auditory pathways. Given the auditory cortex’s relatively limited temporal resolution (Ma et al., 2013), it is likely envelope slope information is initially processed and represented as a spatial map in subcortical nuclei, such as in inferior colliculus, which has better temporal resolution (Khouri et al., 2011). Previous research has demonstrated that primary auditory areas receive well-organized topographic projections from the ventral division of the medial geniculate body (MGv), whereas higher-order areas tend to receive more anatomically and physiologically convergent inputs from MGv (Vasquez-Lopez et al., 2017; Ohga et al., 2018). Even if these envelope steepness maps are topographically projected from MGv to higher-order areas, they may become more heterogeneous due to the convergent projections, similar to how tonotopic maps in higher-order areas become less clear (Issa et al., 2014; Ohga et al., 2018). This transformation of functional organization could render the maps too locally heterogeneous and disordered to be clearly represented with macroscale calcium imaging, thereby obscuring the presence or absence of orthogonal maps in A1 and A2. Nevertheless, it remains possible that envelope information may be represented more sparsely by neurons sharing similar tuning for envelope steepness in a salt-and-pepper manner in higher-order areas. This possibility underscores the need for higher-resolution, single-neuron level mapping to more precisely characterize the functional organization within these regions.

The present study demonstrates orthogonal maps relative to tonotopy, and thus interpretation of the findings may depend on how tonotopic maps are defined and constructed. We visualized tonotopic maps using a straightforward and widely-adopted method of tracking the maximally responding loci at incrementally increasing frequencies (Issa et al., 2014, 2016; Tsukano et al., 2015, 2016; Aponte et al., 2021; Kline et al., 2021; Narayanan et al., 2022). The tonotopic organization and areal parcellation applied in this study have been validated anatomically and histologically in previous works (Horie et al., 2013, 2015; Tsukano et al., 2015; Ohga et al., 2018), supporting the reliability of the maps and minimizing the likelihood of major distortions. However, a more recent study employing a highly systematic mapping approach reported more complex forms of tonotopic organization, particularly in the ventral parts of the mouse auditory cortex (Romero et al., 2020), where splitting, multi-directional tonotopy was reported. If the tonotopic organization is indeed more intricate than previously recognized, this may explain why we did not detect orthogonal temporal maps in ventral higher-order areas. This possibility also warrants further investigation at the single-neuron level within ventral higher-order areas.

While experiments in awake animals are increasingly common, the present study was conducted under anesthesia. Anesthetized preparations offer an important advantage: they reduce variability and noise in neural responses, thereby providing stable conditions for mapping functional organization. Indeed, many foundational topographic features—including auditory tonotopy (Tunturi, 1952; Hind, 1953), visual orientation maps (Bonhoeffer and Grinvald, 1991), and retinotopy (Hubel and Wiesel, 1959; Daniel and Whitteridge, 1961)—were first identified under anesthesia and continue to provide a cornerstone for modern system neuroscience. At the same time, the steepness of rise times is an important component contributing to broader perceptual processes, such as the detection of looming or echoic sounds (Ghazanfar et al., 2002). However, anesthesia imposes constraints on interpretation with respect to perception; it is difficult to directly link rise-ramp maps to perceptual phenomena such as looming or echo perception. These perceptual links should be examined in future studies using awake, behaving animals. Nevertheless, the present findings establish a robust framework for investigating the cortical representation of temporal envelope features.

In the field of otolaryngology, it is well recognized that elderly individuals with normal hearing thresholds often struggle with speech perception, even after adjustments in sound pressure levels (Eisenberg et al., 1995; Peters et al., 1998; Summers and Molis, 2004; George et al., 2006). This clinical challenge highlights the importance of temporal processing in the central nervous system beyond the sensitivity of peripheral hearing receptors (Moore et al., 1992; Noordhoek et al., 2001; Bernstein et al., 2013). Studies using mouse models have explored changes in the central auditory system following the onset of hearing loss, reporting qualitative changes in the arrangement of tonotopic maps in the auditory cortex (Yang et al., 2011; Persic et al., 2020; Resnik and Polley, 2021), raising the possibility that functional organization representing temporal features may likewise be degraded in individuals with impaired speech perception. To fully investigate the dynamism of temporal information in speech, electrophysiology will be indispensable for probing sustained-phase dynamics to assess how envelope steepness is encoded within the body of ongoing speech utterances. Notably, the range of rise-steepness maps identified in this study closely matches the range reported to contribute to human sound lateralization (Kunov and Abel, 1981), suggesting that cortical encoding of envelope steepness may support spatial hearing as well as speech comprehension. The mouse, with its unparalleled genetic accessibility and diverse methodological toolkit—including optogenetics, widefield and two-photon imaging, high-density electrophysiology, and anatomical tracing—represents a powerful system for dissecting the neural basis of temporal envelope processing. By leveraging these approaches, future studies will be able to link mesoscale organization with single-neuron dynamics and move forward addressing the mechanisms underlying speech intelligibility. The present work may serve not only as an experimental foundation but also as a catalyst for further investigations that bridge basic auditory neuroscience with clinical challenges in communication disorders.

---

**Figure S1.**
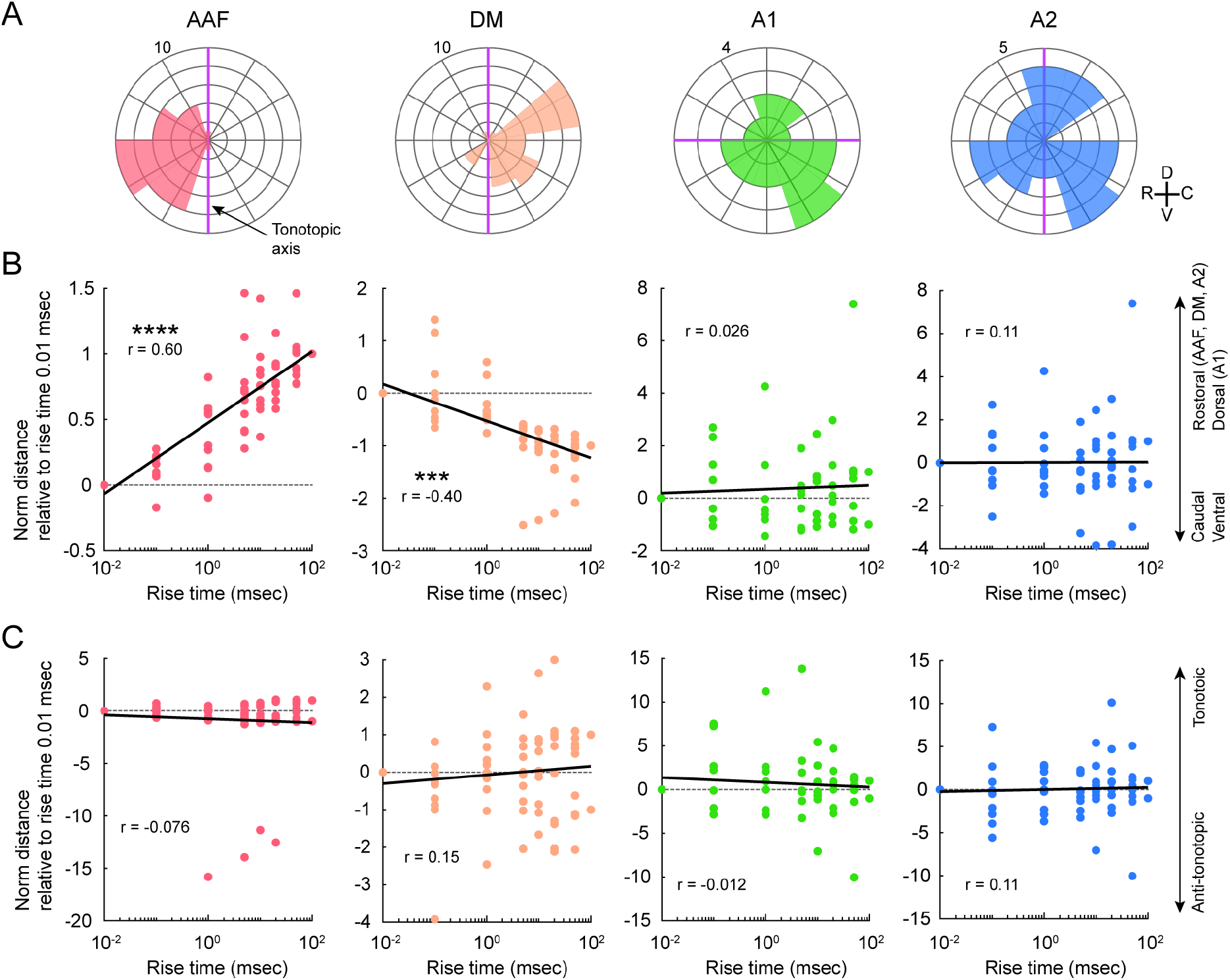
Envelope maps in the left auditory cortex. **(A)** Circular histogram showing the direction of responses of rise-ramp times of 100 msec relative to those for rise-ramp times of 0.01 msec in the left auditory cortical areas. Purple lines indicate the tonotopic axis. Data for 5, 20, 40 kHz are co-plotted. AAF, n = 30 plots; DM, n = 30 plots; A1, n = 26 plots; A2, n = 30 plots from 10 mice. **(B)** Relationship between rise-ramp time and the shift from the peak location for a rise-ramp time of 0.01 msec in the direction orthogonal to tonotopy. AAF, r = 0.60, ****p = 3.9 × 10^-9^, n = 80 plots (Pearson’s correlation); DM, r = -0.40, ***p= 2.1 × 10^-4^, n = 80 plots; A1, r = 0.026, p = 0.83, n = 72 plots; A2, r = 0.11, p = 0.33, n = 80 plots. **(C)** Relationship between rise-ramp time and the shift from the peak location for rise-ramp time of 0.01 msec in the tonotopic direction. AAF, r = -0.076, p = 0.50, n = 80 plots; DM, r = 0.15, p = 0.19, n = 80 plots; A1, r = -0.012, p = 0.92, n = 72 plots; A2, r = 0.11, p = 0.35, n = 80 plots.

**Figure S2.**
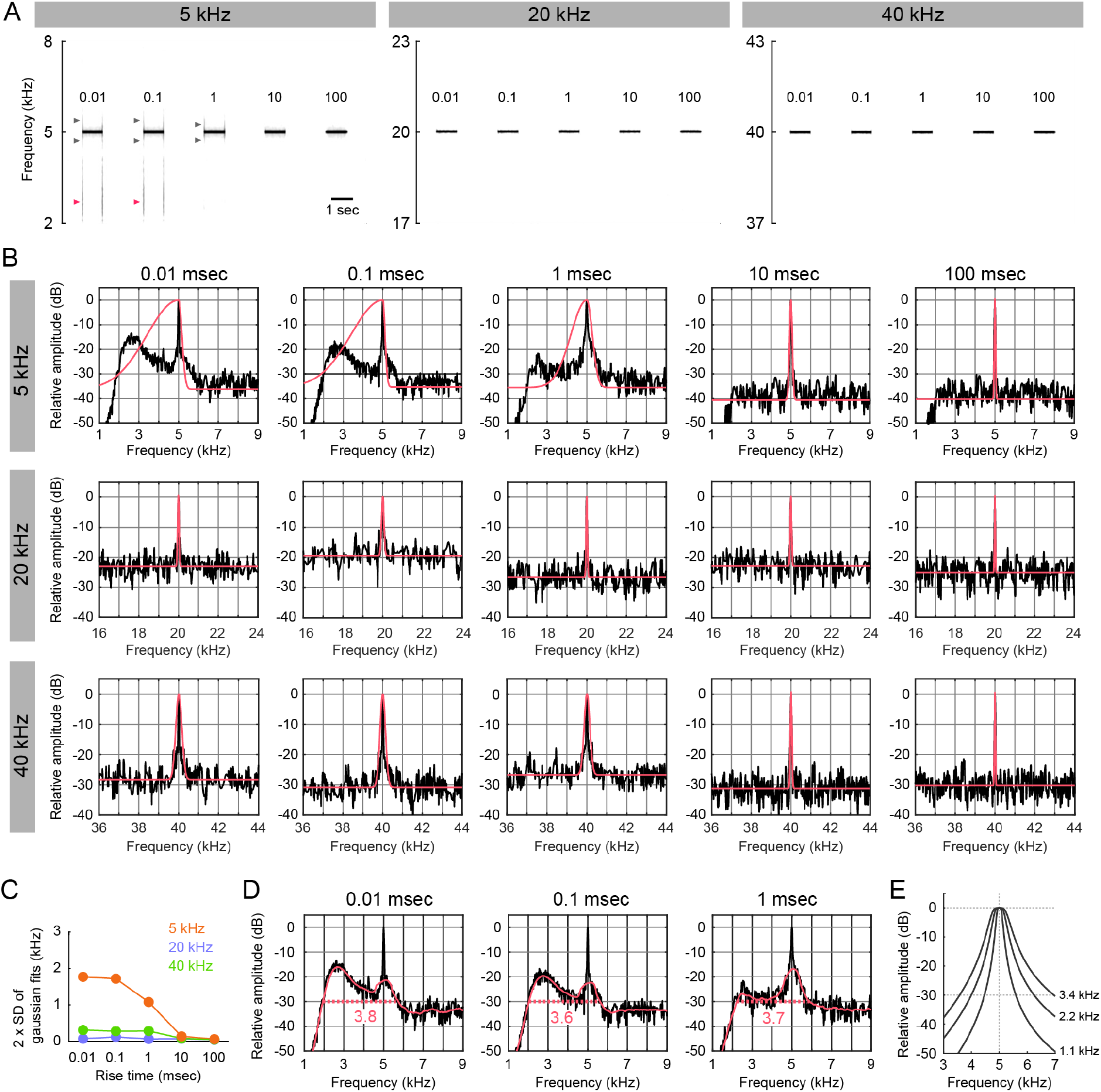
Spectral analysis of pure tones with varying rise times. **(A)** Spectrograms of 5, 20, and 40 kHz pure tones with rise times ranging from 0.01 to 100 ms. Gray arrowheads indicate mild, symmetrical spectral splatter around the carrier frequency, and red arrowheads indicate broader, lower-frequency–biased splatter. **(B)** Amplitude spectra at tone onset across carrier frequencies and rise times. Gaussian fits are shown in red. Y-axis values are expressed in dB relative to the peak amplitude at the carrier frequency. **(C)** Frequency ranges covering ±2 SD of the Gaussian fits shown in (B). **(D)** Additional quantification of splatter width. Red curves represent polynomial fits, and red numbers and dotted lines indicate the width at –30 dB. In (B), Gaussian fits were used to estimate splatter width, but fitting accuracy was poor for 5 kHz tones with rise times of 0.01, 0.1, and 1 msec. Therefore, polynomial fitting was also applied to these data to re-evaluate splatter width, yielding values similar to those obtained with Gaussian fitting shown in Figure 5C. **(E)** Band-pass noise curves added to 5 kHz pure tones. Numbers indicate the width of band-pass noise at –30 dB, matching the frequency range of splatter shown in Figure 5C.

**Figure S3.**
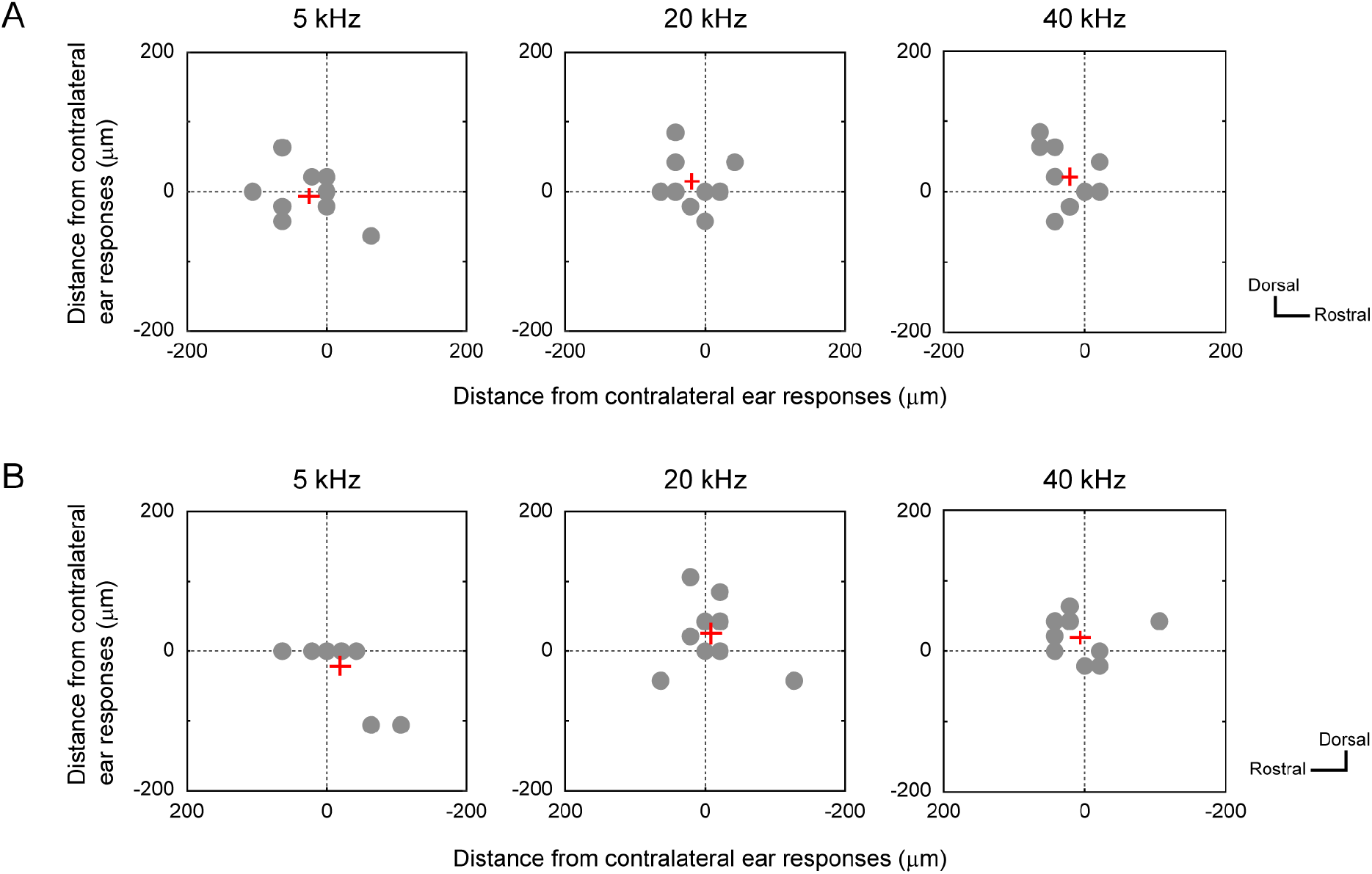
No difference in response locations between contra- and ipsilateral sound inputs. **(A)** Deviation of AAF responses elicited by ipsilateral ear stimulation relative to those elicited by contralateral stimulation in the right hemisphere of the same mice. Results for 5, 20, and 40 kHz at a 10-msec rise time are shown separately. Gray dots indicate each data. Red crosses indicate mean ± SEM. No significant differences were observed along either the horizontal or ventral axis at any frequency (two-sided paired t-test, p > 0.05, n = 9 mice). **(B)** Same as (A), but for AAF responses in the left hemisphere. No significant differences were observed along either axis at any frequency (p > 0.05, n = 9 mice).

**Figure S4.**
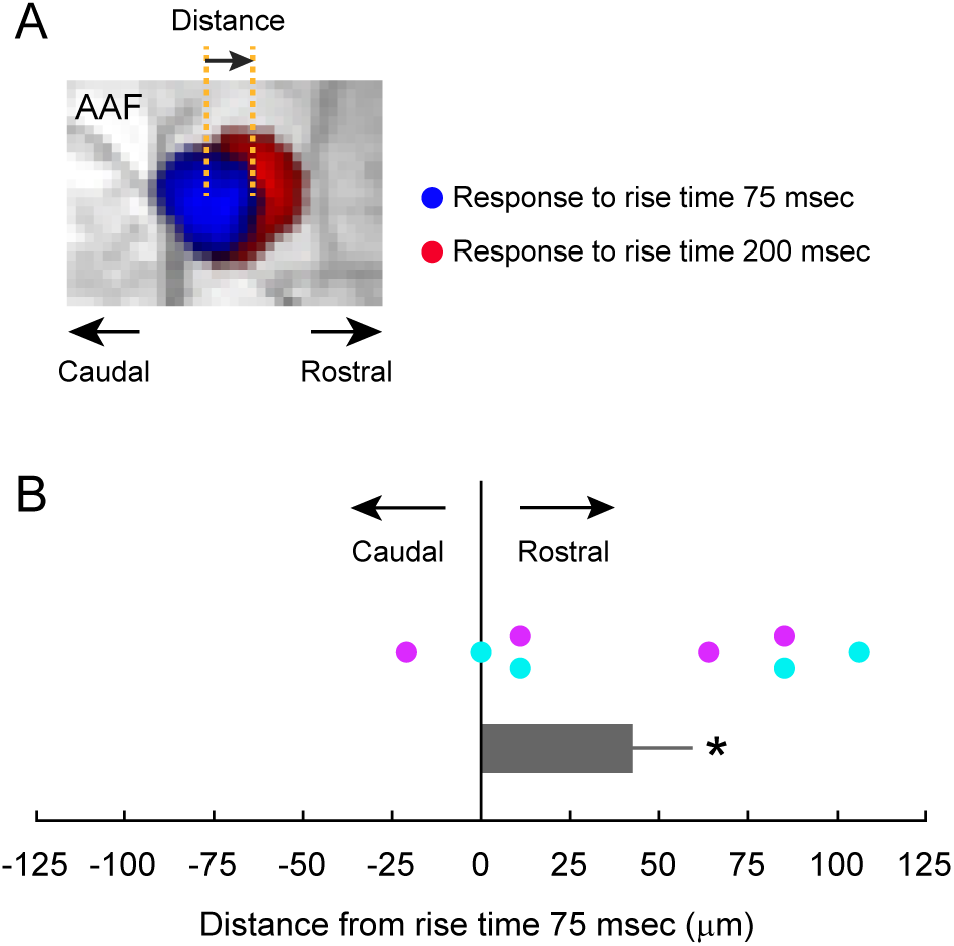
Envelope maps during the ramping phase. **(A)** Explanation of analysis. Positional shifts of 5 kHz responses are evaluated between rise times of 75 and 200 msec. **(B)** Plots of rostro-caudal deviation between the peak responses for a rise-ramp time of 200 msec relative to that for a rise-ramp time of 75 msec in AAF. Positive values indicate that the response for a rise-ramp time of 200 msec is rostral to that for 75 msec, while negative values indicate it is caudal. Data for AAF in the right (cyan) and left hemisphere (purple) are co-plotted (n = 8 plots from 4 mice). Data are shown in mean ± SEM. *p = 0.039 (two-sided paired t-test).

## Methods

### Animals

The experimental procedures in the present study were approved by the Committee for Animal Care at Niigata University. All the experiments were performed in accordance with the approved guidelines and regulations. We used 5–7-wk-old Thy1-GCaMP6f mice (C57BL/6J-Tg(Thy1-GCaMP6f)GP5.5Dkim/J) acquired from Jackson Laboratories. The animals were housed in cages with *ad libitum* access to food pellets and water and kept on a 12-h light/dark cycle.

### Sound stimuli

Sound waveforms were made using an application (RPvdsEx) of a sound generator (System3, Tucker-Davis Technologies, Alachua, FL) at a sampling rate of 130 kHz. Waveforms were low-pass filtered at 100 kHz (3625, NF, Kanagawa, Japan). The auditory stimuli were transmitted to both ear canals simultaneously, except in the experiments shown in Figure S3. Each ear canal received sound through a dedicated sound guide tube, which was connected to a speaker (EC1, Tucker-Davis Technologies). The sound intensity was set at ∼60 dB SPL at the location of the ear canal. Sound intensity was calibrated using a sound level meter (NA-42S, RION, Kokubunji, Japan) with a microphone and pre-amplifier (UC54 and NH-17, RION). To evaluate map shifts related to onset ramp steepness, 1-sec tones were presented with rise/fall times ranging 0.01–200 msec across 8 conditions (0.01, 0.1, 1, 5, 10, 20, 50, and 100 msec; Figures 2–4, S1) or 2 conditions (75 and 200 msec; Figure S2). To assess the effect of sound intensity, 5-kHz tones with a fixed 0.01-ms rise time were presented at 30–60 dB SPL. To test the effect of broadband frequency noise, Gaussian noise with bandwidths of 0, 1.1, 2.2, and 3.4 kHz (–30 dB relative to the peak) was added to 5-kHz tones with a 100-ms rise time. The noise profiles were generated using a 2nd-order Butterworth filter with cutoff frequencies of 0.1, 0.2, or 0.3 kHz and are shown in Figure S2. To visualize tonotopy in Figure 2, we used 1-sec tones with amplitude modulation at 20 Hz.

### Sound recording and spectral analysis

A condenser microphone with a pre-amplifier (UC54 and NH-17, RION), which provides a flat frequency response up to 200 kHz, was connected to the tubing from a speaker (EC1). Sound waveforms were digitized at a 192-kHz sampling rate. To evaluate spectral splatter associated with changes in onset ramp steepness, time–frequency analysis was performed using short-time Fourier transform (STFT). Spectrograms were computed in MATLAB (MathWorks) with a Hanning window (42.7-ms temporal resolution, 11.7-Hz frequency resolution), 75% overlap, and 2^14^-point FFT. Alterations of these parameters within reasonable ranges did not influence the outcome of the analysis. The STFT output was converted to dB as:

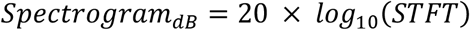

Frequency vectors at the onset were extracted, and amplitude distributions were extracted from STFT. The amplitude spectrum y was fitted with either a Gaussian or an asymmetric Gaussian model. For the symmetric case, the model was

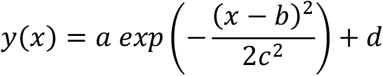

where *x* is the input frequency, *a* is the amplitude, *b* is the peak position, *c* is the standard deviation (*σ*), and *d* is the baseline offset. For conditions where the spectrum was skewed (e.g., 5 kHz tones with very short rise times), an asymmetric Gaussian was used:

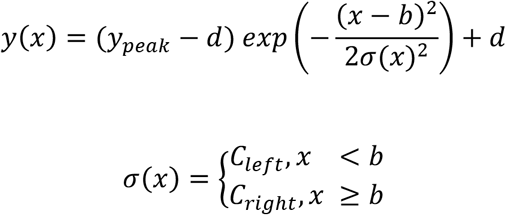

From the fit, we calculated the standard deviations (*σ*_*left*_, *σ*_*right*_) and the bandwidths at –30 dB attenuation level, and defied *σ* as the average of *σ*_*left*_ and *σ*_*right*_. For each condition, goodness-of-fit was assessed using the coefficient of determination R^2^. Fits with R^2^<0.7 were regarded as low quality and re-fitted with adjusted initial parameters.

When a polynomial function was fitted, weighted least squares was used:

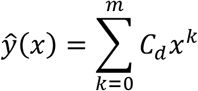

where *m* is the polynomial degree and *C*_*k*_ are coefficients. Weights were assigned to emphasize the region around the spectral peak and deemphasize baseline regions. The quality of the fit was evaluated by the coefficient of determination R^2^.

### Surgery

Mice were deeply anesthetized using urethane (1.65 g/kg, i.p.; Wako, Osaka, Japan) and administered atropine (0.5 mg/kg, s.c.; Terumo, Tokyo, Japan). After local anesthesia using 1% Xylocaine (Aspen Pharmacare, Durban, South Africa), the skin and temporal muscle over the auditory cortex in both hemispheres were incised, leaving the skull intact. A piece of metal was attached to the skull with dental resin, and the head was fixed by screwing the metal piece onto a manipulator. The exposed surface of the skull was covered with liquid paraffin (Wako) to keep the skull transparent during imaging. The rectal temperature was maintained at 37°C during the surgery using a heat pad throughout experiments.

### Calcium imaging

About 30 min after the completion of the surgery, macroscale calcium imaging was performed transcranially to reveal functional maps in the auditory cortex. Bilateral cortical images (128 × 168 pixels) were recorded using a cooled CCD camera system (ORCA-R2, Hamamatsu Photonics, Hamamatsu, Japan) attached to a microscope (M165FC, Leica Microsystems), which was controlled with a LabVIEW-based system, RatioImaging Recorder (E.I.SOL, Minato-ku, Japan). GCaMP6f was excited by LED light (λ = 460 nm; UHP-Mic-LED-460, Opto-Line, Warabi, Japan), and its fluorescence was detected through band-bass filters (λ = 500–550 nm). The area covered by one pixel was 21.2 × 21.2 µm. Images were taken at the sampling rate of 20 (Figures 3–5, S1–3) or 50 Hz (Figure S4).

Image analyses were performed using custom-made MATLAB codes (MathWorks, Natick, MA). Images of the same stimuli were averaged over 15 trials. These images were converted to ΔF/F_0_, with the baseline intensity (F_0_) obtained by averaging the intensity values during the prestimulus period of 250 msec. A spatial averaging of 5 × 5 pixels was applied. To obtain response images, consecutive image frames over a 0–400 ms window following stimulus onset were collected, and the mean values across this time window were calculated for each pixel (Figures 3–5, S1–3). For the analysis of Figure S4, the frame corresponding to 40–60 msec following tonal onset was selected. The locations of pixels with the maximum responses were identified within each auditory area in a semi-automated manner: Reference pixels were put within each auditory areas according to the overall spatial pattern, and pixels with the largest ΔF/F_0_ value were automatically found within a radius of 8 pixels from the reference point. When multiple pixels were identified as peak pixels with identical values, the averaged coordinate across these pixels was considered the peak location.

For the analysis presented in Figures 3–5, S1–3, a coordinate matrix of [3 frequencies × 8 rise steepness × 10 mice] was obtained for each auditory area. For each mice, the coordinate matrix of [3 frequencies × 8 rise steepness] was rotated so that the tonotopic direction between 5 kHz and 40 kHz (AAF, A1, and A2) or 20 kHz and 40 kHz (DM) with a rise time of 0.01 msec was aligned vertical (AAF, DM, and A2) or horizontal (A1). When creating two-dimensional plots, the distance between 5 kHz and 40 kHz (AAF, A1, and A2) or 20 kHz and 40 kHz (DM) was normalized to 1 to average across mice. After averaging coordinate data across frequencies, angles of each data point relative to the tonotopic axis were plotted in a circular histogram, with the data distributed across 10 bins. In plots showing the relationships between rise time and distance, the distance between the peak locations for rise times of 0.01 and 100 ms was set to 1 for each frequency for normalization. NaN was put in data points when clear peak location of auditory areas was not identified. For the analysis of Figure S4, the rostrocaudal difference in the peak locations for 5 kHz between rise times of 75 msec and 200 msec was measured.

To provide the temporal profile of GCaMP signals in Figure 2C, the intensity values within a 5 × 5 pixel area centered around the peak location of the response were averaged. To present representative responses for each frequency in Figure 2D, response images were color-coded and thresholded using RatioImaging Recorder. To generate the merged image shown in Figure 2E, responsive pixels were extracted from each auditory area. Within each area, responses were normalized to the local peak, and threshold levels (0.70–0.85% of the maximum peak) were determined by visual inspection using RatioImaging Recorder to best reveal the tonotopic organization. Pixels within responsive regions were identified using a flood-fill algorithm, initiated from seed points placed in activated areas. The extracted peak response regions were then color-coded and merged into a single image using ImageJ (NIH, Bethesda, MD). For Figure 2G, response images were separately normalized for each auditory area and thresholded to extract peak response regions, yielding binary images. Threshold levels (0.70–0.85% of the maximum peak) were similarly chosen by visual inspection to optimally visualize the envelope steepness organization. Responsive regions were extracted using the same flood-fill algorithm and merged using Illustrator (Adobe, Sun Jose, CA).

### Statistics

Statistical analyses were conducted using MATLAB. Correlations were evaluated using Pearson’s correlation. The two-sided paired t-test was used to evaluate whether data distribution was significantly different than zero. Only p-values less than 0.05 were highlighted by asterisks.

## Author Contributions

K.T. and H.T. conceptualized the project. K.T., T.G., T.Y., and S.O. performed experiments. K.T., T.G., T.Y., S.O., and H.T. analyzed the data. K.T. and H.T. wrote the manuscript. K.T., H.T., and A.H. revised the manuscript. All the authors approved the final version of the manuscript.

## Acknowledgements

This work was supported by JSPS KAKENHI No. 20K09750 (T.K.). We thank Michellee M. Garcia at the University of North Carolina at Chapel Hill for her helpful comments and for editing the English in this manuscript.

## Data and codes availability

Data and codes supporting this study are available from corresponding authors upon reasonable requests.

